# Glutamine metabolism regulates Th2 cell differentiation via the α-ketoglutalate-dependent demethylation of histone H3K27

**DOI:** 10.1101/184648

**Authors:** Makoto Kuwahara, Maya Izumoto, Hiroaki Honda, Kazuki Inoue, Yuuki Imai, Junpei Suzuki, Saho Maruyama, Masaki Yasukawa, Masakatsu Yamashita

## Abstract

The acquisition of T cell functions seems to be closely linked to the reprogramming of the metabolic pathway. However, the impact of metabolic changes on the differentiation of helper T cell subsets remains unclear. We found that TCR-mediated activation of glutamine metabolism regulates Th2 cell differentiation via the supplementation of α-ketogulutalate (α-KG) and histone H3K27 demethylation. Deprivation of glutamine or pharmacological inhibition of glutamine metabolism blocks the induction of Th2 cell differentiation without affecting Stat6 phosphorylation. The methylation status of H3K27 at the Th2 cytokine gene locus was significantly increased in Th2 cells cultured under glutamine-deprived conditions. The inhibitory effect of glutamine-deprivation was antagonized by α-KG, and the α-KG-dependent induction of Th2 cell differentiation was reduced in *utx-* and *jmjd3-deficient* naïve CD4 T cells. These findings show that the glutamine-α-ketoglutarate axis is crucial to regulating the epigenetic status at the Th2 cytokine gene locus and subsequent Th2 cell differentiation.

## INTRODUCTION

On recognition of a foreign antigen, naïve CD4 T cells are rapidly activated, expanding and differentiating into various helper T (Th) cell subsets, including Th1, Th2 and Th17 cells, which produce different cytokines and have distinct effector functions (1, 2). Naïve CD4 T cells can also differentiate into inducible regulatory T (iTreg) cells, which suppress undesired immune responses. The excessive activation of Th cells or misdirected Th cell differentiation results in the promotion of inflammatory disorders, such as allergic inflammation and autoimmune diseases (3, 4).

Emerging evidence suggests that T cells dramatically alter their metabolic activity during T cell receptor (TCR)-mediated activation (5-7). This change in the metabolic status is termed “metabolic reprogramming” and plays an important role in the regulation of the T cell-mediated immune response. Similar to other non-proliferating cells, naïve T cells use fatty acid oxidization and/or a low rate of glycolysis and subsequently oxidize glucose-derived pyruvate via oxidative phosphorylation (OXOPHOS) to generate ATP (8, 9). Upon activation, T cells immediately shift their metabolic program to anabolic growth and biomass accumulation that support the rapid expansion of the cells and the acquisition of an effector function. Both effector CD4 and CD8 T cells have high rates of glucose consumption and lactate production despite the availability of sufficient oxygen to oxidize glucose completely (aerobic glycolysis), similar to aggressive tumors (8), known as the Warburg effect (10). In this process, pyruvate derived from glucose is converted into lactate. In addition, pyruvate can be transported into mitochondria, where it is converted to acetyl-CoA.

In rapidly proliferating cells including activating T cells, several kinds of amino acids are used as fuel for OXPHOS. The nonessential amino acid glutamine is the most abundant amino acid in the blood and serves as a source of carbon and nitrogen for the synthesis of proteins, lipids and amino acids (11). Proliferating cells import extracellular glutamine and catabolize it via glutaminolysis in both the cytosol and mitochondria (12). Glutaminolysis consists of two steps (13). In the first step, glutaminase converts glutamine to glutamate. In the second step, glutamate dehydrogenase and aspartate aminotransferases convert glutamate into the tricarboxylic acid (TCA) cycle intermediate α-ketoglutarate (α-KG), which is consumed through OXPHOS or a reductive TCA cycle (13) (14). Glutamine metabolism also supplements the pyruvate pool, which is predominantly formed from glucose. As a consequence of the rapid metabolism of glutamine, the rapidly proliferating cells secrete lactate, alanine and NH4+ (11). The effector T cells also rapidly take up glutamine, and glutamine is required for the maximum cell growth and proliferation of T cells (15, 16).

The important role of metabolic reprograming in epigenetic regulation has been established (17). Metabolites act as essential cofactors in the epigenetic regulation of transcription. Acetyl-CoA and NAD+ are involved in the regulation of histone acetylation. Acetylation of histones in achieved via the action of histone acetyl transferases (HATs), which attach an acetyl group to lysine residues using acetyl-CoA as a substrate. Sirtuin family histone deacetylases are dependent on a readily available pool of NAD+ for their deacetylase activity. The α-KG regulates the enzymatic activity of Tet methylcytosine dioxygenase 2 (Tet2) and JumonjiC (JmjC) family histone demethylases (18). It has been well established that changes in histone modifications (acetylation, methylation, etc.) of the cytokine gene loci play an important role in regulating differentiation of the Th cell subset. However, the relationship between metabolic reprograming and the epigenetic regulation of Th cell differentiation is not completely understood.

Therefore, in this study, we assessed the intrinsic role of glutamine metabolism in epigenetic regulation of Th cell differentiation.

## Results

### IL-4 accelerates glutaminolysis and glycolysis in activated CD4 T cells

Proliferating T cells import extracellular glutamine and catabolize it via glutaminolysis. The intracellular level of glutamate was dramatically increased by anti-TCR-β plus anti-CD28 mAb stimulation, and upregulation of intracellular glutamate was completely dependent on extracellular L-glutamine *in vitro* (Fig. 1A). The intracellular glutamate level peaked at 24 h after the TCR-stimulation and gradually decreased (Figs. 1A and 1B). We found that the intracellular glutamate level was reduced by IL-4 (Fig. 1B), indicating that IL-4 affects amino acid metabolism that of glutamine, in activated CD4 T cells.

**Figure 1.**
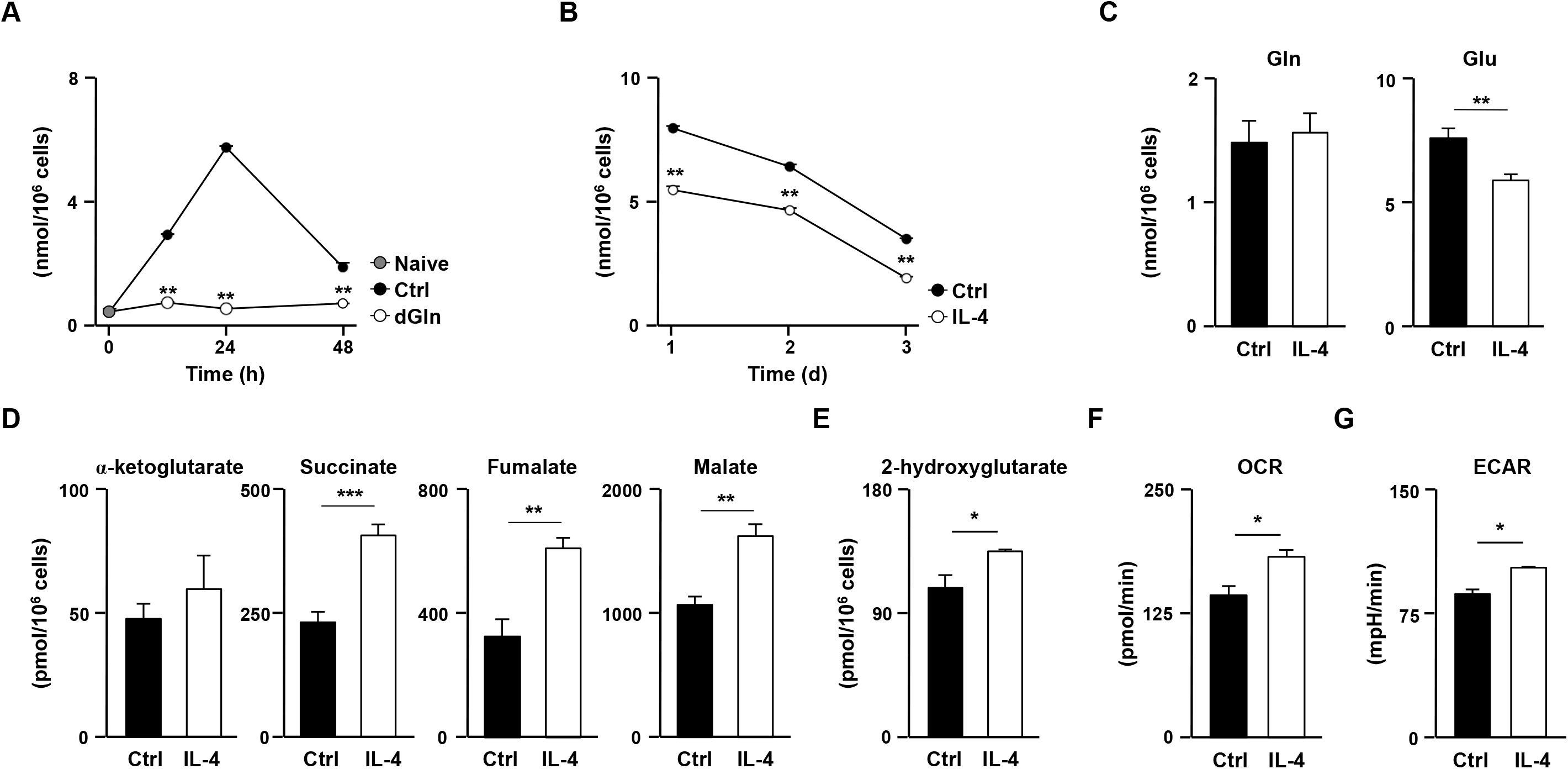
IL-4 accelerates glutamine metabolism. (**A**) Naïve CD4 T cells were stimulated with anti-TCR-β mAb, anti-CD28 mAb and IL-2 under control (Ctrl) or glutamine-deprived (dGln) conditions for the indicated durations. The intracellular glutamate concentrations are shown with the standard deviations (n = 3: biological replicates). **P<0.01 (Student’s t-test). (**B**) The effect of IL-4 on intracellular glutamate concentration in activated CD4 T cells. Naïve CD4 T cells were stimulated as in (**A**) in the presence or absence of IL-4 for 3 days. The intracellular glutamate concentrations are shown with the standard deviation (n = 3: biological replicates). **P<0.01 (Student’s t-test). (**C**) The intracellular concentrations of glutamine and glutamate in activated CD4 T cells cultured in the presence (IL-4) or absence (Anti-IL-4) of IL-4. Naïve CD4 T cells were stimulated with anti-TCR-β/anti-CD28 mAb in the presence of IL-2 and anti-IFN-γ mAb for 48 h. (D, E) The intracellular concentrations of metabolic intermediates in the TCA cycles (**D**) and 2-hydroxyglutarate, a metabolite of α-KG (**E**) in activated CD4 T cells cultured as in (**C**). The concentrations are shown with the standard deviation (n = 3: biological replicates). *P<0.05, **P<0.01, ***P<0.001 (Welch’s t-test). (**F**) The basal oxygen consumption rate (OCR) and extracellular acidification rate (ECAR) in the activated CD4 T cells cultured as in (**C**). The results are shown with the standard deviation (n = 3: biological replicates). *P<0.05 (Student’s *t*-test).

To assess the effect of IL-4 on the metabolic status in activated CD4 T cells comprehensively, naïve CD4 T cells were stimulated with an anti-TCR-β mAb plus anti-CD28 mAb for 48 h in the presence or absence of IL-4 and subjected to metabolic profiling of 116 metabolites. The intracellular level of glutamate was significantly reduced by IL-4, whereas the level of glutamine remained unchanged (Fig. 1C). The concentrations of metabolic intermediates of the TCA cycle such as α-KG, succinate, fumarate and malate, increased in IL-4-treated cells (Fig. 1D). The level of 2-hydroxyglutarate, a metabolite ofα-KG, was also increased by IL-4 administration (Fig. 1E). In contrast, the levels of citrate, cis-aconitate and isocitrate were unaffected by IL-4 treatment (Fig. 1-Figure Supplement 1A). Furthermore, the oxygen consumption rate (OCR) was also augmented by IL-4 treatment (Fig. 1F) implying accelerated OXOPHOS via glutaminolysis in IL-4-treated activated CD4 T cells. In addition, the intracellular concentrations of glycolytic metabolites such as glucose 6-phosphate, fructose 6-phosphate and fructose 1,6-diphosphate were decreased in IL-4-treated activated CD4 T cells, whereas 3-and 2-phosphoglycerate, pyruvate and lactate were increased (Fig. 1-Figure Supplement 1B). The extracellular acidification rate was also enhanced by IL-4 (Fig. 1G). These data indicate that both glutaminolysis and anaerobic glycolysis are facilitated in activated CD4 T cells by IL-4.

### Glutamine metabolism is required for Th2 cell differentiation

We next focused on the role of glutamine metabolism in Th2 cell differentiation since IL-4 plays an important role in the induction of Th2 cells. The increase in intracellular glutamate by TCR-stimulation was abolished by the deprivation of extracellular L-glutamine (Fig. 1A); we therefore cultured naïve CD4 T cells under Th2 conditions with L-glutamine-deprivation medium to investigate the role of glutaminolysis in Th2 cell differentiation. The L-glutamine level in this culture medium is reduced not completely depleted, because a substantial amount of L-glutamine is present in fetal calf serum (complete medium: 3mM L-Gln, glutamine-deprived medium: 0.15mM Gln).

The generation of IL-4-, IL-5-and IL13-producing cells was substantially decreased in naïve CD4 T cells cultured in the L-glutamine-deprivation medium (Fig. 2A). A reduction in Th2 cytokine production was also detected in Th2 cells induced under L-glutamine-deprived conditions (Fig. 2B). The glutamine deprivation reduced the acetylation levels of histone H3K27 (Fig. 2C) and H3K9 at the Th2 cytokine gene locus (Fig. 2-Figure Supplement 1A). In sharp contrast, the H3K27 tri-methylation level was increased (Fig. 2D). Unexpectedly, we found that the level of H3K4 tri-methylation at the Th2 cytokine gene loci did not decrease, but rather increased, in the Th2 cells differentiated under the glutamine-deprived conditions (Fig. 2-Figure Supplement 1B). The glutamine deprivation did not affect the IL-4-induced tyrosine phosphorylation of Stat6 (Fig. 2E, or the IL-4-dependent induction of Gata3 (Fig. 2F). These results indicate that glutamine deprivation affects the histone modification status without affecting the Stat6/Gata3 pathway, and thereby inhibiting Th2 cell differentiation.

**Figure 2.**
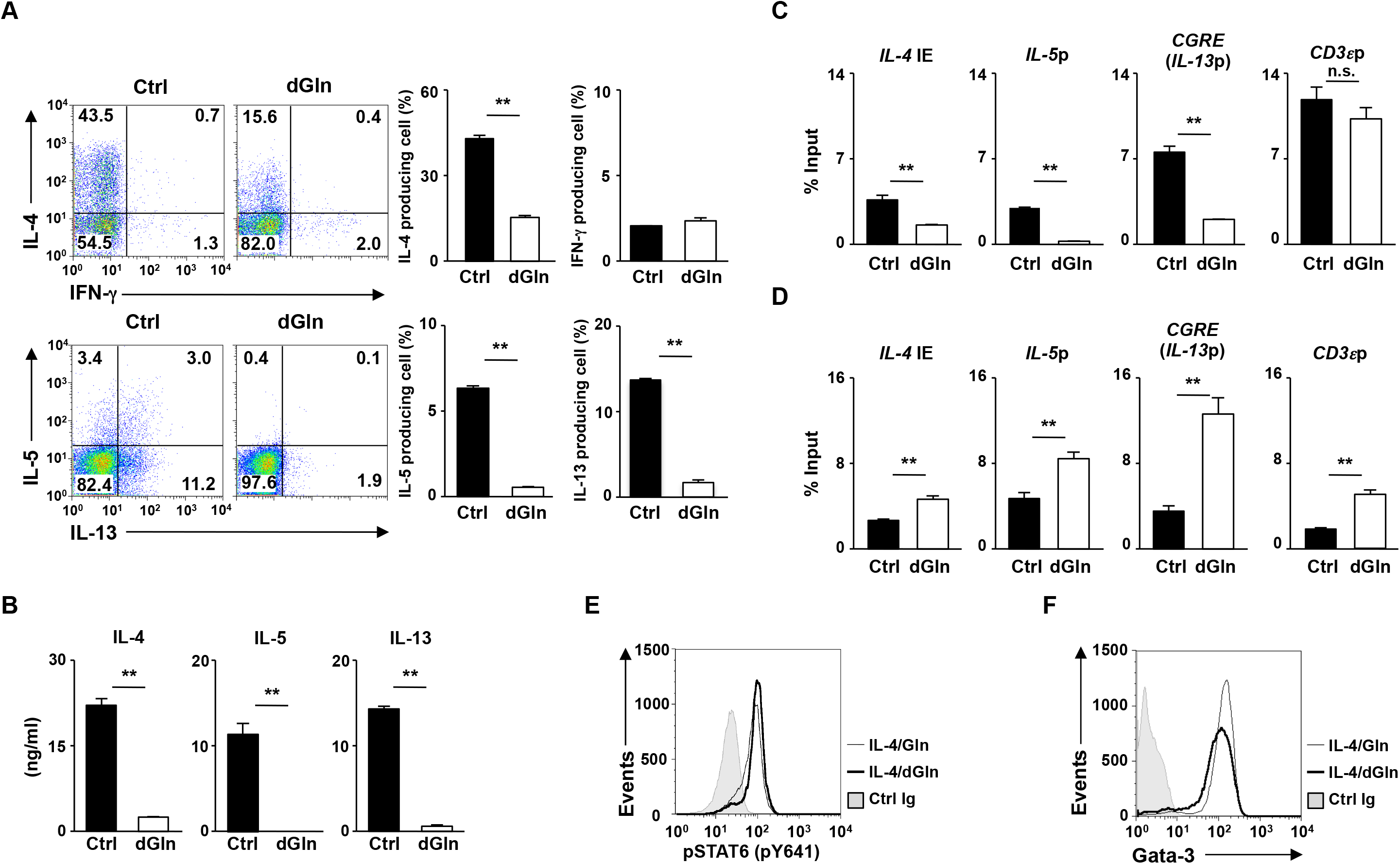
Glutamine is required for efficient Th2 differentiation. (**A**) The results of the intracellular FACS analysis of IFN-γ/IL-4 (upper panel) and IL-13/IL-5 (lower panel) in the naïve CD4 T cells culture under normal (Ctrl) or glutamine-deprived (dGln) Th2 conditions for 5 days. The percentages of cells are indicated in each quadrant. The percentages of IL-4-, IL-5, IL-13 and IFN-γ-producing cells in three independent cultures are shown with the standard deviation (right panel). **P< 0.01 (Student’s t-test). (**B**) The results of the ELISA for IL-4, IL-5 and IL-13 in the supernatants of the cells in (**A**) restimulated with an immobilized anti-TCR-β mAb for 16 h are shown with the standard deviation (n = 3: biological replicates). **P<0.01 (Student’s t-test). (**C and D**) The results of the ChIP assay with a quantitative PCR analysis of the histone modification (H3K27ac and H3K27me3) status at the Th2 cytokine gene and CD3ε gene loci in the Th2 cells induced under L-glutamine-deprived Th2 conditions (dGln); the results are presented relative to those of input DNA with the standard deviations (n = 3: technical replicates). **P< 0.01, n.s.: not significant (Student’s t-test). (**E**) The IL-4-dependent tyrosine phosphorylation of Stat6 in naïve CD4 T cells cultured under L-glutamine-deprived Th2 conditions (dGln) for 2 h. (**F**) The results of the intracellular FACS analysis of Gata3 in Th2 cells stimulated under L-glutamine-deprived Th2 conditions (dGln) for two days. The data are representative of at least three independent experiments with similar results.

To confirm the role of glutamine metabolism in Th2 cell differentiation, we next assessed the effect of amino oxyacetic acid (AOA), an inhibitor of transaminase on Th2 cell differentiation. The treatment of naïve CD4 T cells cultured under Th2 conditions with AOA during TCR stimulation substantially reduced the generation of IL-4-, IL-5-and IL-13-producing cells (Fig. 3A). A significant number of IFN-γ-producing CD4 T cells were generated in the AOA-treated Th2 cultures (Fig. 3A). The AOA-treated Th2 cells produced smaller amounts of Th2 cytokines than did the untreated control CD4 T cells (Fig. 3B). The AOA treatment also reduced the acetylation levels of histone H3K27 (Fig. 3C) and H3K9 at the Th2 cytokine gene locus (Fig. 3-Figure Supplement 1A). Enhanced H3K27 and H3K4 tri-methylation levels were detected in the AOA-treated Th2 cells (Fig. 3D and Fig. 3-Figure Supplement 1B). The IL-4-induced tyrosine phosphorylation of Stat6 was unaffected by the AOA treatment (Fig. 3E). The IL-4-dependent induction of Gata3 in activated CD4 T cells was moderately inhibited by the AOA treatment (Fig. 3F). These results demonstrate a critical role for glutamine metabolism in regulating the chromatin status at the Th2 cytokine gene locus and subsequent Th2 cell differentiation.

**Figure 3.**
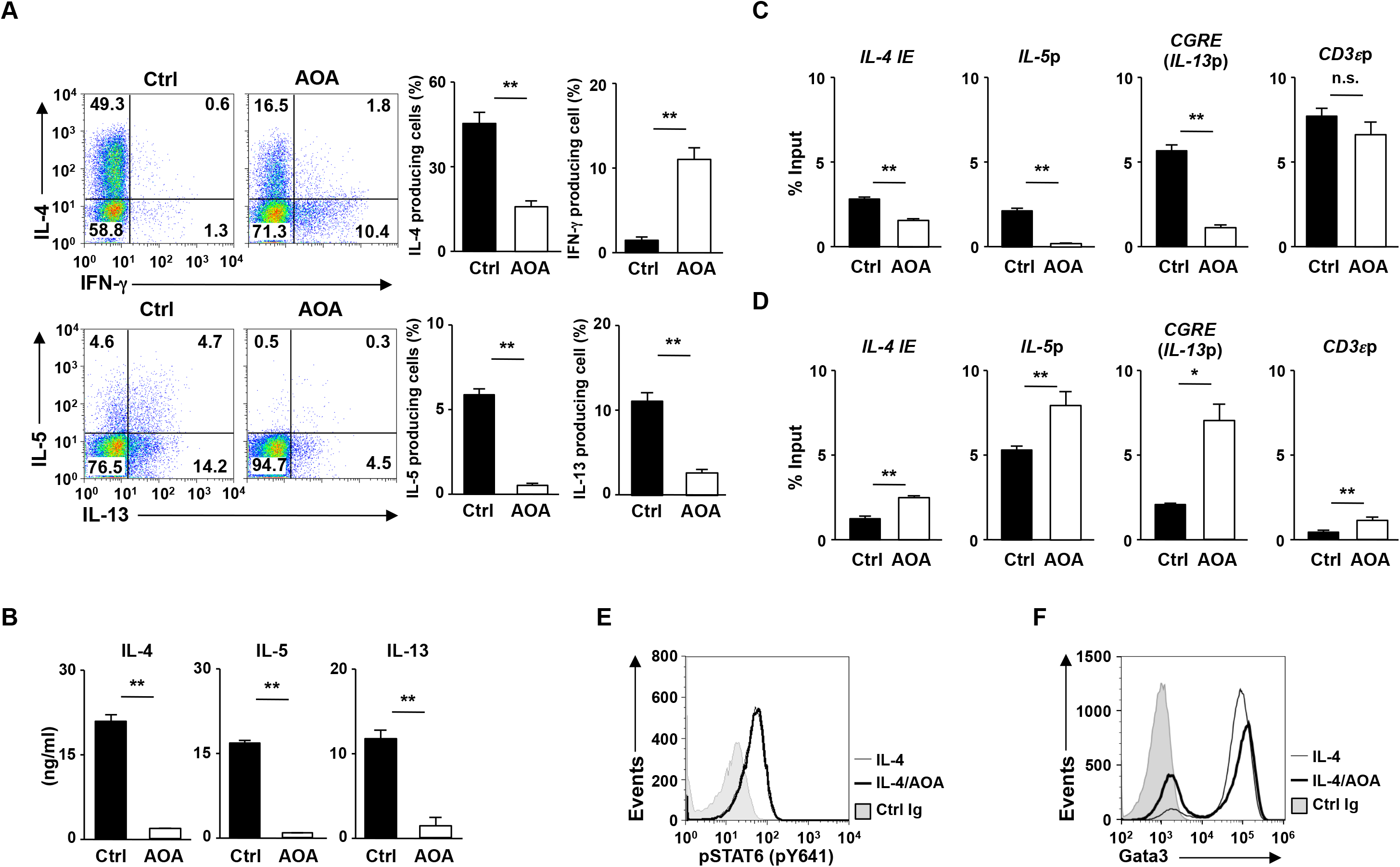
The pharmacological inhibition of glutamine metabolism suppressed Th2 differentiation. (**A**) The results of the intracellular FACS analysis of IFN-γ/IL-4 (upper) and IL-13/IL-5 (lower). Naïve CD4 T cells were cultured under Th2 conditions in the presence or absence of aminooxyacetic acid (AOA: 1 mM) for two days. The percentages of IL-4-and IFN-γ-producing cells in three independent cultures are shown with the standard deviations (right) (n = 3: biological replicates). **P< 0.01 (Student’s t-test). (**B**) The results of an ELISA of cytokines in the supernatants of the cells in (**A**) restimulated with immobilized anti-TCR-β for 16 h. The averages of three independent cultures are shown with the standard deviations (n = 3: biological replicates). **P<0.01, n.s.: not significant (Student’s t-test). (C and D) The results of the ChIP assay with a quantitative PCR analysis of the histone modification (H3K27ac and H3K27me3) status at the Th2 cytokine gene and CD3ε gene loci in the Th2 cells induced as in (**A**); the results are presented relative to those of the input DNA with the standard deviations (n = 3: technical replicates). **P<0.01, n.s.: not significant (Student’s t-test). (**E**) The IL-4-dependent tyrosine phosphorylation of Stat6 in naïve CD4 T cells cultured under Th2 conditions in the presence or absence of AOA for 4 h. (**F**) The results of the intracellular FACS analysis of Gata3 in naïve CD4 T cells cultured under Th2 conditions in the presence or absence of AOA for two days. The data are representative of at least three independent experiments with similar results.

### The glutamine-α-KG axis regulates Th2 cell differentiation

We next wanted to determine the molecular mechanisms by which glutamine metabolism regulates Th2 cell differentiation. α-KG, a metabolite of glutamate, regulates the enzymatic activity of JmjC family histone demethylases, Tet2 and PDH hydroxylases, suggesting that glutaminolysis is involved in the cellular differentiation processes (18). An increased degree of histone H3K27 methylation at the Th2 cytokine gene locus was detected in Th2 cells cultured under glutamine-deprived conditions (Fig. 2E). We therefore assessed the effect of α-KG administration on Th2 cells cultured under glutamine-deprived conditions. The generation of IL-4-, IL-5-and IL-13-producing Th2 cells under glutamine-deprived conditions was significantly enhanced by α-KG administration (Fig. 4A). The increased production of Th2 cytokines (IL-4, IL-5 and IL-13) by α-KG administration was also detected by an enzyme-linked immunosorbent assay (ELISA) (Fig. 4B). Furthermore, the level of histone H3K27 tri-methylation of the Th2 cytokine gene locus was significantly reduced by α-KG (Fig. 4C). In contrast, the reduced cell division of developing Th2 cells under glutamine-deprived conditions showed only partial restoration by α-KG (Fig. 4D). These results suggest that glutamine metabolism regulates Th2 cell differentiation via the supplementation of α-KG.

**Figure 4.**
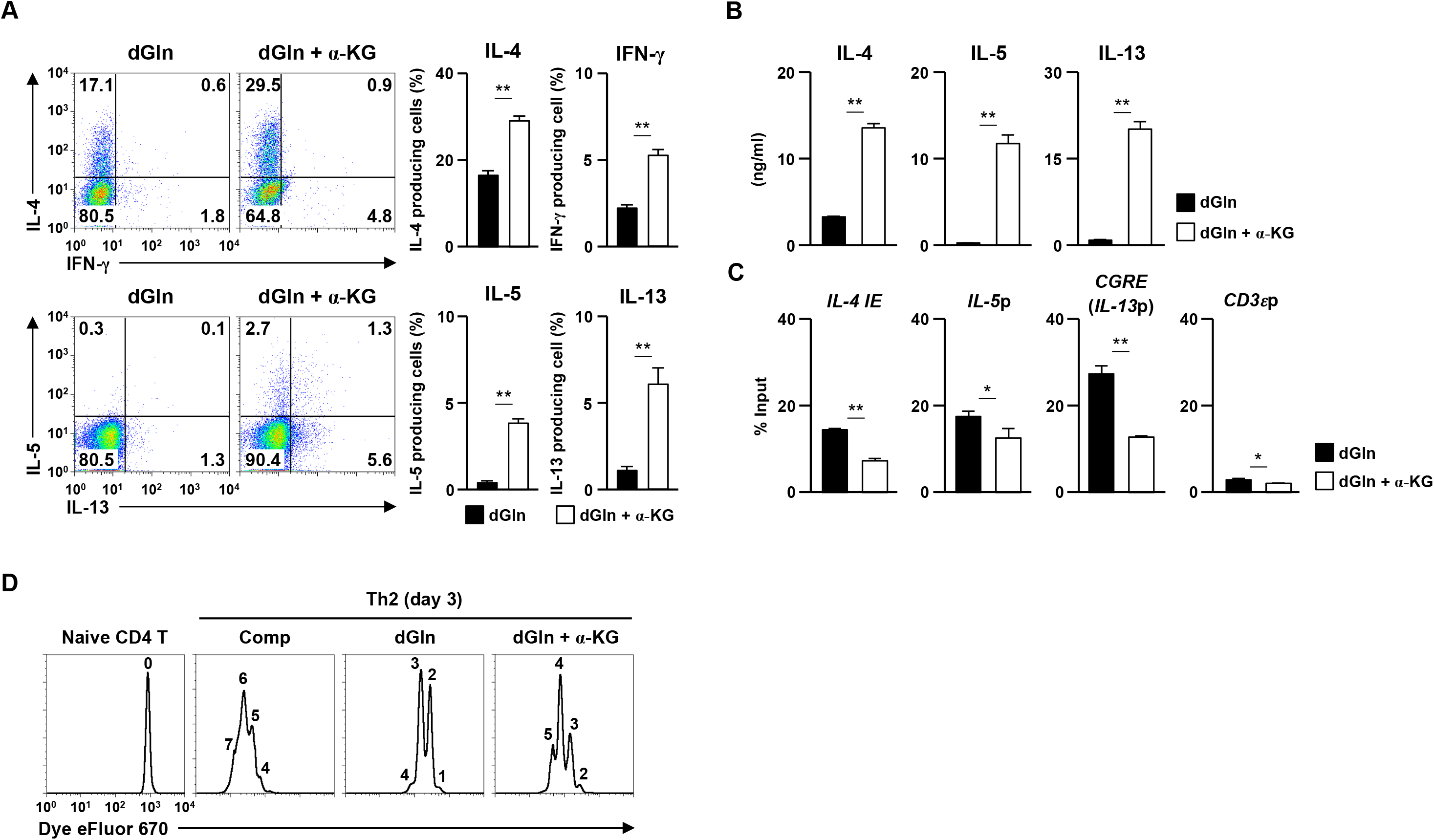
The α-KG-dependent demethylation of the Th2 cytokine gene loci is required for the efficient Th2 differentiation. (**A**) The results of the intracellular FACS analysis of IL-4/IFN-γ (upper) and IL-5/IL-13 (lower) in naïve CD4 T cells cultured under glutamine-deprived Th2 conditions (IL-4, 1 ng/ml) in the absence (dGln) or presence (dGln + α-KG) of cell-permeating dimethyl-α-ketoglutarate (DM-α-KG, 2 mM). Naïve CD4 T cells were cultured under glutamine-deprived Th2 conditions in the absence or presence of DM-α-KG for two days, and then the cells were further expanded with IL-2 plus IL-4 for three days with or without DM-α-KG. The numbers of cells are indicated in each quadrant. The percentages of IL-4-, IL-5-, IL-13-, and IFN-γ-producing cells in three independent cultures are shown with the standard deviations (right). (**B**) Results of the ELISA of cytokines in the supernatants of the cells in (**A**) restimulated with an immobilized anti-TCR-β mAb for 16 h. The averages of three independent cultures are shown with the standard deviations (n = 3: biological replicates). **P<0.01 (Student’s t-test). (**C**) The levels of histone H3K27 tri-methylation at the Th2 cytokine and CD3ε gene loci in naïve CD4 T cells cultured under glutamine-deprived Th2 conditions in the presence (α-KG) or absence (dGln) of DM-α-KG for two days. The results are presented relative to those of the input DNA with the standard deviations (n = 3: technical replicates). *P<0.05, **P<0.01 (Student’s *t*-test) (**D**) A representative FACS analysis of glutamine-dependent proliferative responses of CD4 T cells. Naïve CD4 T cells were labeled with Dye eFluor 670 and cultured under normal (Ctrl) or glutamine-deprived Th2 conditions in the presence (dGln + α-KG) or absence (dGln) of DM-α-KG for three days. The data are representative of at least three independent experiments with similar results.

### Histone H3K27 demethylases utx and jmjd3 is required for efficient Th2 cell differentiation

Two kinds of histone H3K27me2/me3 demethylases have been identified: Kdm6a (Utx) and Kdm6b (Jmjd3) (19, 20). We therefore established T cell-specific utx-deficient (uřx^flox flox^ x CD4-Cre Tg) and *jmjd3*-deficient (*jmjd3^ũoxlũox^* x CD4-Cre Tg) mice to assess the role of H3K27 demethylation in α-KG-dependent Th2 cell differentiation. The bindings of the Utx and Jmjd3 at the regulatory regions of the Th2 cytokine gene locus were detected by a ChIP-qPCR analysis (Fig. 5A). In the absence of *utx,* the generation of IL-4-, IL-5-and IL-13-producing cells moderately decreased (Fig. 5B). Decreased production of IL-4, IL-5 and IL-13 in uřx-deficient Th2 cells was confirmed by an ELISA (Fig. 5C). The reduced generation of IL-5-and IL-13-producing cells was also observed in *jmjd3*'-deficient naïve CD4 T cell culture, whereas the number of IL-4-produicing cells remained unaffected (Fig. 5B). In contrast, the production of IL-4, IL-5 and IL-13 was significantly decreased by jmjd3 deficiency (Fig. 5C). The tri-methylation level of histone H3K27 at the Th2 cytokine gene locus was increased in both *utx-* and *jmjd3-deficient* Th2 cells (Fig. 5D). Finally, we assessed the α-KG-dependent acquisitions of IL-5 and IL-13 production ability by *utx-* and jmjd3-deficient CD4 T cells. The α-KG-dependent induction of IL-5 and IL-13 production was reduced in both *utx-* and jmjd3-deficient CD4 T cells (Fig. 5E). In contrast, the decreased α-KG-dependent induction of IL-4 production was only detected in utx-deficient CD4 T cells, not in jmjd3-deficient cells.

**Figure 5.**
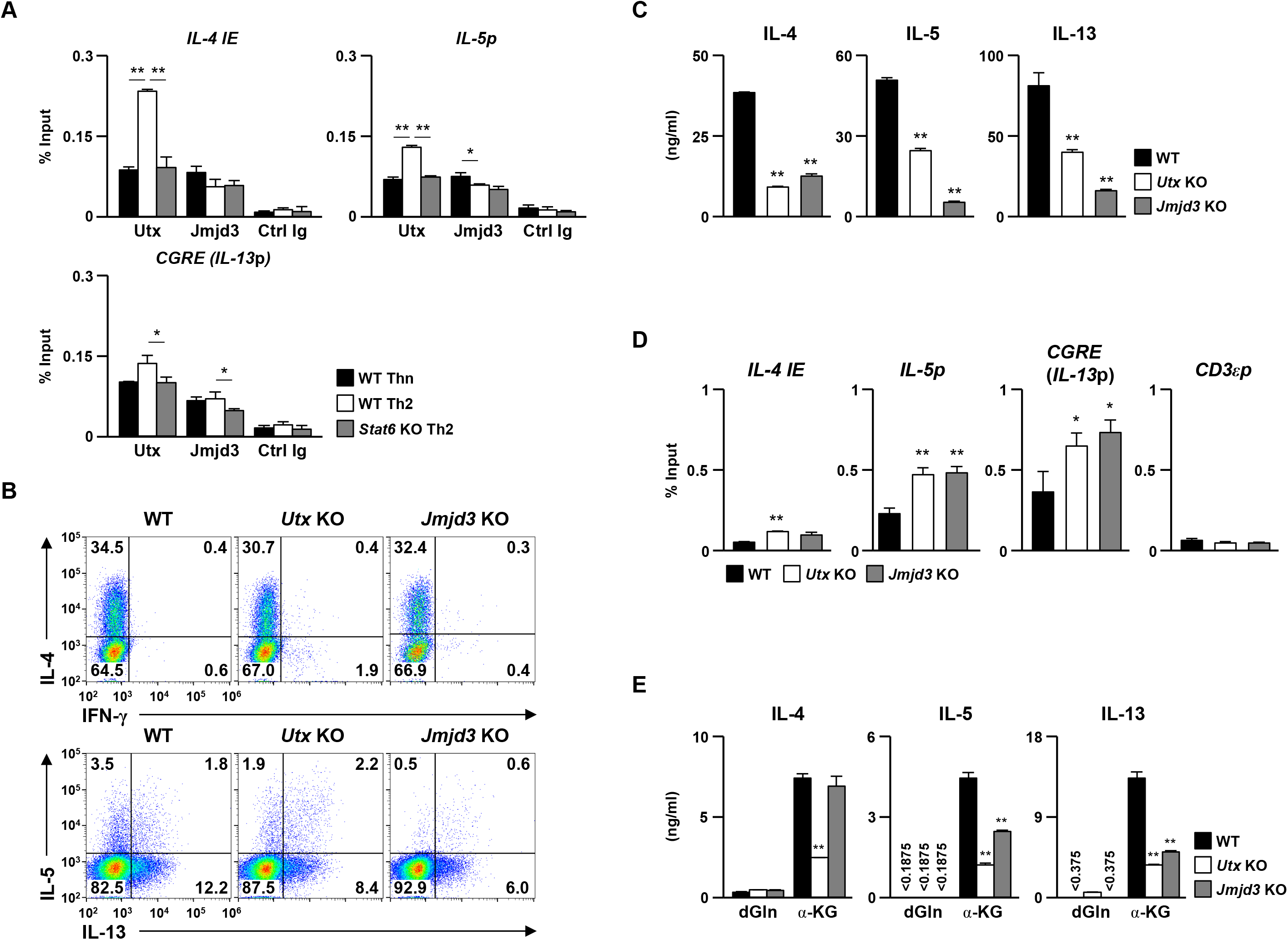
Histone H3K27 demethylases Utx and Jmjd3 are required for the Th2 differentiation. (**A**) The bindings of Utx and Jmjd3 at the Th2 cytokine gene locus were determined by ChIP-qPCR. Naïve CD4 T cells were cultured under Th2 conditions for 5 days and subjected to a ChIP assay. The results are presented relative to those of the input DNA with the standard deviations (n = 3: technical replicates). *P<0.05, **P< 0.01 (Student’s t-test). (**B**) Representative results of the intracellular FACS analysis of IL-4/IFN-γ (upper) and IL-5/IL-13 (lower) in the wild-type (WT), Utx-deficient (*Utx* KO) and *Jmjd3-deficient (Jmjd3* KO) naïve CD4 T cells cultured under Th2 conditions. The numbers of cells are indicated in each quadrant. (**C**) Results of the ELISA of cytokines in the supernatants of the cells in (**B**) restimulated with an immobilized anti-TCR-β mAb for 16 hours. The averages of three independent cultures are shown with the standard deviations (n = 3: biological replicates). **P<0.01 (Student’s t-test). (**D**) The levels of histone H3K27 tri-methylation (upper) and acetylation (lower) at the Th2 cytokine and CD3ε gene loci in the cells in (**A**). The results are presented relative to those of the input DNA with the standard deviations (n = 3: technical replicates). *P<0.05, **P<0.01 (Student’s t-test). (**e**) Results of the ELISA of Th2 cytokines in the supernatant of WT, *Utx* KO, *Jmjd3* KO Th2 cells restimulated with an immobilized anti-TCR-β mAb for 16 h. WT, *Utx* KO, and *Jmjd3* KO naïve CD4 T cells were cultured under glutamine-deprived Th2 conditions in the presence (α-KG) or absence (dGln) of DM-α-KG for five days. The averages of three independent cultures are shown with the standard deviations (n = 3: biological replicates). **P<0.01 (Student’s t-test).

## Discussion

We herein showed that the inhibition of glutamine metabolism or deprivation of extracellular glutamine reduced Th2 cell differentiation *in vitro.* Glutamine metabolism is accompanied by increases in the production of α-KG. α-KG regulates the enzymatic activity of Tet2, JmjC family histone demethylases and PDH hydroxylases (18). It is well known that the differentiation of Th2 cells is regulated by histone and DNA methylation (21-23). We showed that H3K27me3, an inhibitory histone mark, at the Th2 cytokine gene locus was increased in naïve CD4 T cells cultured under AOA-treated or glutamine-deprived Th2 conditions. Therefore, glutamine metabolism may regulate the differentiation of Th2 cells by regulating the histone H3K27 methylation level at the Th2 cytokine gene locus. We found that the glutamine-α-KG axis regulates the H3K27 methylation status at the Th2 cytokine gene locus. We also showed that the histone H3K27 demethylases, Utx and Jmjd3 are involved in the Th2 cell differentiation. The demethylation of histone H3K27 seems to be induced in a Stat6/Gata3-independent manner. We and other authors have previously reported the Stat6-and Gata3-dependent induction of histone H3K9 acetylation at the Th2 cytokine gene locus in developing Th2 cells (24, 25). Thus, the glutamine-α-KG axis and the Stat6-Gata3 pathway cooperate to induce Th2 cell differentiation.

The expressions of several possible glutamine transporters (Slc1A5, Slc3A2, Slc7A5, and Slc38A1) were induced in CD4 T cells by TCR stimulation (16). It was previously reported that Slc1a5 (Asct2), an amino acids transporter, facilitates the glutamine uptake and controls inflammatory CD4 T cell responses by regulating Th1 and Th17 differentiation but not Th2 cell differentiation (26). Asct2 appears to regulate Th1 and Th17 cell differentiation by modulating the mTOR-signaling pathway. L-glutamine is required for the transport of essential amino acids including L-leucine into cells via Slc7a5 (LAT1/CD98) (27), and cytoplasmic L-leucine activates the mTOR-signaling pathway (28, 29). The critical role of mTORC1 signaling in Th1 and Th17, but not Th2 differentiation has been previously demonstrated (30, 31). The Asct2-dependent incorporation of glutamine may be required for the amino acid-dependent activation of mTORC1 in CD4 T cells. Although the important role of Asct2 in Th1 and Th17 cell differentiation was clearly demonstrated in an analysis of *Asct2*-deficient CD4 T cells, the role of the glutamine metabolism was not assessed. In the present study, we showed that glutamine metabolism is required for Th2 differentiation. Furthermore, we found that the pharmacological inhibition of glutamine metabolism or deprivation of glutamine resulted in the impaired generation of IL-17A-producing CD4 T cells but not the impaired generation of IFN-γ-producing cells. Glutamine metabolism is accompanied by the increased production of α-KG. We demonstrated that α-KG-dependent demethylation of histone H3K27 at the Th2 cytokine gene locus is involved in Th2 differentiation. In addition, the differentiation of Th17 is also regulated by α-KG-dependent histone H3K27 demethylation (M.K. & M.Y. unpublished observation). Therefore, it is possible that glutamine metabolism controls Th2 and Th17 differentiation via the supplementation of α-KG. Thus, we conclude that glutamine regulates Th cell differentiation and subsequent CD4 T cell-dependent immune response through multiple pathways.

We detected the TCR-mediated upregulation of intracellular glutamate in activated CD4 T cells, and the increase in the intracellular glutamate concentration was completely dependent on extracellular glutamine. These results suggest that extracellular glutamine is transported into activated CD4 T cells and converted into glutamate, presumably by glutaminolysis. Glutaminolysis consists of two steps (13). In the first step, glutaminases (Gls1 and Gls2) and/or aspartate synthase (Asns) converts glutamine to glutamate. In the second step, glutamate dehydrogenase (Glud1), alanine aminotransferases (Alt1 and Alt2) and aspartate aminotransferases (Ast1 and Ast2) convert glutamate into α-KG (13) (14). In addition, glutamate is also converted into α-KG by phospho-serine aminotransferase 1 (Psat1) during *de novo* serine synthesis (32). Serine is used as a source of purine biosynthesis via supplementation of glycine (33, 34). A metabolic profiling analysis indicated that IL-4 accelerates purine synthesis in activated CD4 T cells (M.K. & M.Y. unpublished observation). Therefore, it is possible that IL-4 may augments the intracellular α-KG level by facilitating the serine biosynthesis pathway.

## Materials and Methods

### Mice

*Cre* TG mice under the control of the *Cd4* promoter were purchased from Jackson Laboratory. Stat6-deficient mice were kindly provided by Dr. Shizuo Akira (Osaka University). *Utx^flox/flox^* mice were established by Drs. Kazuki Inoue and Yuuki Imai (Ehime University). *Jmjd3^flox/flox^* mice were kindly provided by Dr. Hiroaki Honda (Hiroshima University). C57BL/6 mice were purchased from Clea (Clea Japan, Inc., Tokyo, Japan). Gene-manipulated mice with C57BL/6 background were used in all experiments. All experiments using mice received approval from the University Administrative Panel for Animal Care. All animal care was conducted in accordance with the guidelines of Ehime University.

### Reagents and antibodies

L-glutamine-free RPMI1640 (Cat#183-2165) and aminooxyacetic acid (Cat#037-06191) were purchased from Wako Pure Chemical Industries (Osaka, Japan). Dimethyl 2-oxoglutarate (Dimethyl α-ketoglutarate; DM-α-KG) was obtained from Tokyo Chemical Industries (cat#K0013; Tokyo, Japan). The antibodies used intracellular staining as follow: anti-IL-4-phycoerythrin (PE) mAb (11B11; BD Bioscience, San Jose, CA, USA), anti-IFN-γ-FITC mAb (XMG1.2; BD Bioscience), anti-IL-5-allophycocyanin (APC) (TRFK5; BioLegend, San Diego, CA, USA), anti-IL-13-PE (eBio13A; eBioscience, San Diego, CA, USA), anti-Stat6 (pY641)-Alexa fluor 647 (J71-773.58.11; BD bioscience), and anti-Gata3-Alexa Fluor 647 (L50-823; BD Biosciences). The antibodies used for the ChIP assay were as follows: anti-Utx pAb (cat#3938; abcam, Cambridge, UK), anti-Jmjd3 pAb (cat#85392; abcam), anti-histone H3K4me3 pAb (cat#39159; Active Motif, Carlsbad, CA, USA), anti-histone H3K27ac pAb (cat#39133; Active Motif) and anti-histone H3K27me3 pAb (cat#39155; Active Motif). All antibodies were diluted and used according to the manufacturer’s instructions.

### CD4 T cells stimulation and differentiation in vitro

Naïve CD4 T (CD44^low^CD62L^high^CD25^negative^) cells were prepared using a CD4+CD62L+ T cell isolation kit II (cat#130-093-227; Miltenyi Biotec, San Diego, CA, USA). Naïve CD4 T cells (1.5×10^6^ cells) were stimulated with an immobilized anti-TCR-β mAb (3 ¼g/ml, H57-597; BioLegend) and an anti-CD28 mAb (1 ¼g/ml, 37.5; BioLegend) for 2 days under the conditions indicated. The cells were then transferred to a new plate and further cultured in the presence of cytokines. The cytokine conditions were as follows: Neutral (Thn) conditions, IL-2 (10 ng/ml), anti-IL-4 mAb (5 ¼g/ml, 11B11; BioLegend), and anti-IFN-γ mAb (5 ¼g/ml, R4-6A2; BioLegend); Th2 conditions, IL-2 (10 ng/ml), IL-4 (1 ng/ml, 3 ng/ml), and anti-IFN-γ mAb (5 ¼g/ml, R4-6A2; BioLegend).

### Intracellular staining of cytokines and transcription factors

For the intracellular staining of cytokines, cells were differentiated in vitro and stimulated with an immobilized anti-TCR-β mAb (3 μg/ml, H57-597; BioLegend) for 6 hours with monensin (2 μM, cat#M5273; Sigma-Aldrich), and intracellular staining was then performed as described previously (35). For the intracellular staining of phospho-stat6, the cells tested without restimulation were stained using a BD phosflow Lysis/Fix buffer (cat#558049; BD Bioscience) and BD phosflow Perm Buffer III (cat#558050; BD Bioscience) according to the manufacturer’s protocol. For the intracellular staining of Gata3, the cells tested without restimulation were stained using a Transcription Factor Staining Buffer Kit according to the manufacturer’s protocol (cat#TNB-0607-KIT; TONBO biosciences, San Diego, CA, USA). Flow cytometry (FACS) was performed using a FACSCalibur (BD Biosciences) and a Galios (Beckman Coulter) instruments, and the results were analyzed using the FlowJo software program (Tree Star, Ashland, OR, USA).

### ELISA assay

Cells were restimulated with an immobilized anti-TCR-β mAb (10 μg/ml) for 16 hours. The concentrations of IL-4, IL-5 and IFN-γ in the supernatants were determined using ELISAs, as described previously (35). The concentrations of IL-13 in the supernatants were determined using the DuoSet ELISA Kit (cat#DY413; R&D systems, Minneapolis, MN, USA).

### Quantitative reverse transcriptase polymerase chain reaction (qRT-PCR)

Total RNA was isolated using TRIzol reagent and cDNA was synthesized using the Superscript VILO cDNA synthesis kit (cat#11754; Thermo Fisher Scientific, Waltham,

MA, USA). Quantitative RT-PCR was performed as described previously (35) using a Step One Plus Real-Time PCR Systems (Life Technologies).

### ChIP assay

The Magna ChIP kits (cat#MAGNA0001 and #MAGNA0002; Merck Millipore, Billerica, MA, USA) were used for the ChIP assay according to the manufacturer’s protocol.

## Cell proliferation assay

Naïve CD4 T cells were labeled with Cell Proliferation Dye eFlour 670 (Cat#65-0840; Invitrogen: Thermo Fisher Scientific, 5 μM) according to the manufacturer’s protocol. The cells were stimulated with anti-TCR-β mAb plus anti-CD28 mAb under Th2 conditions for two days, and then further expanded under Th2 conditions.

### Metabolic profiling

Metabolome measurements and data processing were performed through a facility service at Human Metabolome Technology Inc. (Yamagata, Japan). In brief, naïve CD4 T cells were stimulated with anti-TCR-β mAb plus anti-CD28 under Thn or Th2 conditions for 48 hours. The cells (3×10^6^ cells) were washed with 5% (w/w) mannitol and then lysed with 800 μl of methanol and 500 μl of Milli-Q water containing internal standards (H3304-1002, Human Metabolome Technology Inc.) and left to rest for another 30 sec. The extract was obtained and centrifuged at 2,300 × *g* at 4 °C for 5 min, and then 800 μl of the upper aqueous layer was centrifugally filtered through a Millipore 5-kDa cutoff filter at 9,100 × *g* at 4 °C for 120 min to remove proteins. The filterate was centrifugally concentrated and re-suspended in 50 μl of Milli-Q water for the capillary electrophoresis-mass spectrometry (CE-MS). Cationic compounds were evaluated using the in the positive mode of capillary electrophoresis-time of flight-mass spectrometry (CE-TOFMS) and anionic compounds were evaluated using the in the positive and negative modes of capillary electrophoresis-tandem mass spectrometry (CE-MS/MS) in accordance with methods developed by Soga et al.

### Intracellular glutamate colorimetric assay

The concentration of intracellular glutamate in naïve and activated CD4 T cells cultured under Thn and Th2 conditions was determined using a Glutamate Colorimetric Assay Kit in accordance with the manufacture’s instructions (cat#K629-100; BioVision, Milpitas, CA, USA).

### ChIP-qPCR primers

The specific primers and Roche Universal probes used in ChIP-qPCR were as follows: IL-4 IE: 5’ CCCAAAGGAGGTGCTTTTATC 3’ (forward), 5’ AAATCCGAAACTGAGGAGTGC 3’ (reverse), probe #75; CGRE: 5’ CTCTCCTGGTGGCGTGTT 3’ (forward), 5’ CTTTGCGCACCCTTGAAC 3’ (reverse), probe #53; IL-5p: 5’ TCACTTTATCAGGAATTGAGTTTAACA 3’ (forward), 5’ GATCGGCTTTTCTTGAGCAC 3’ (reverse), CD3sp: 5’ ACACTTCCTGTGTGGGGTTC 3’ (forward), 5’ CTGAAGAAGGCACCAGACG 3’ (reverse), probe #16

## AUTHOR CONTRIBUTIONS

M.K. performed the experiments, analyzed the data and wrote the manuscript; M.I., J.S., S.M. and M.Y. performed and supported the experiments; H.H., K. I., and Y.I. established and provided the *Jmjd3^flox/flox^* and *Utx^flox/flox^* mice. M.Y. conceptualized the research, directed the study and edited the manuscript.

## ACKNOWLEDGEMENTS

We thank A. Tamai her excellent technical assistance. This work was supported by the JSPS KAKENHI Grant Numbers 816K191580, 917H040860, 25860376, the Mochida Memorial Foundation for Medical and Pharmaceutical Research, the Takeda Science Foundation, the Uehara Memorial Foundation and the Naito Foundation.

## Figures-Figure supplements legends

**Figure 1-Figure Supplement 1.**
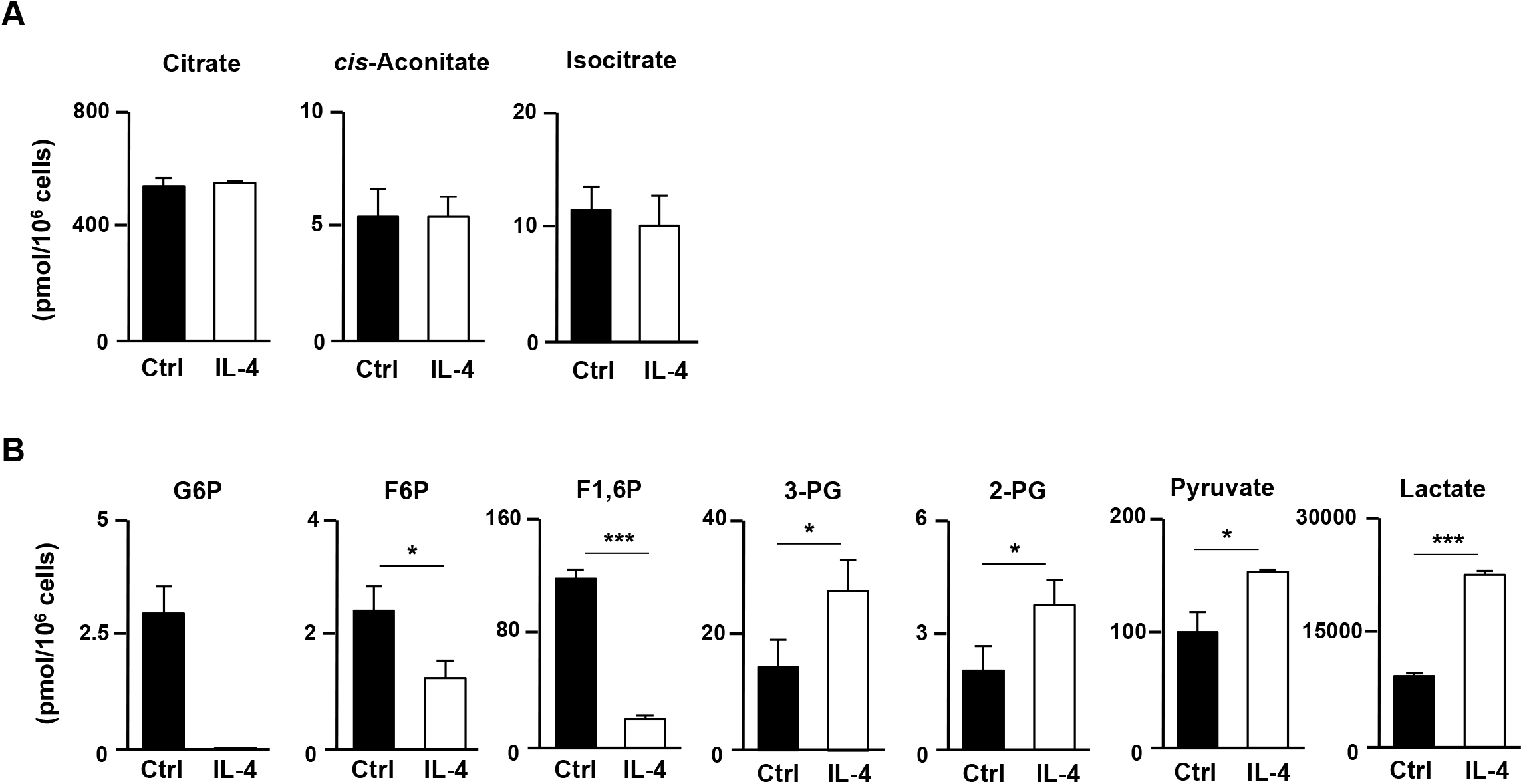
IL-4-dependent metabolic alterations in activated CD4 T cells. The results of intracellular concentrations of metabolic intermediates in the TCA cycle (**A**) or the glucose metabolism (**B**) in naïve CD4 T cells stimulated in the presence of anti-IL-4 mAb (Ctrl) or IL-4 conditions (IL-4). WT naïve CD4 T cells were stimulated with anti-TCR-β, anti-CD28 mAb, IL-2 and anti-IFN-γ mAb in the presence of anti-IL-4 mAb or IL-4 for 48 h. The results are shown with the standard deviations (n = 3: biological replicates). *P<0.05, **P<0.01, ***P<0.001 (Welch’s t-test).

**Figure 2-Figure Supplement 1.**
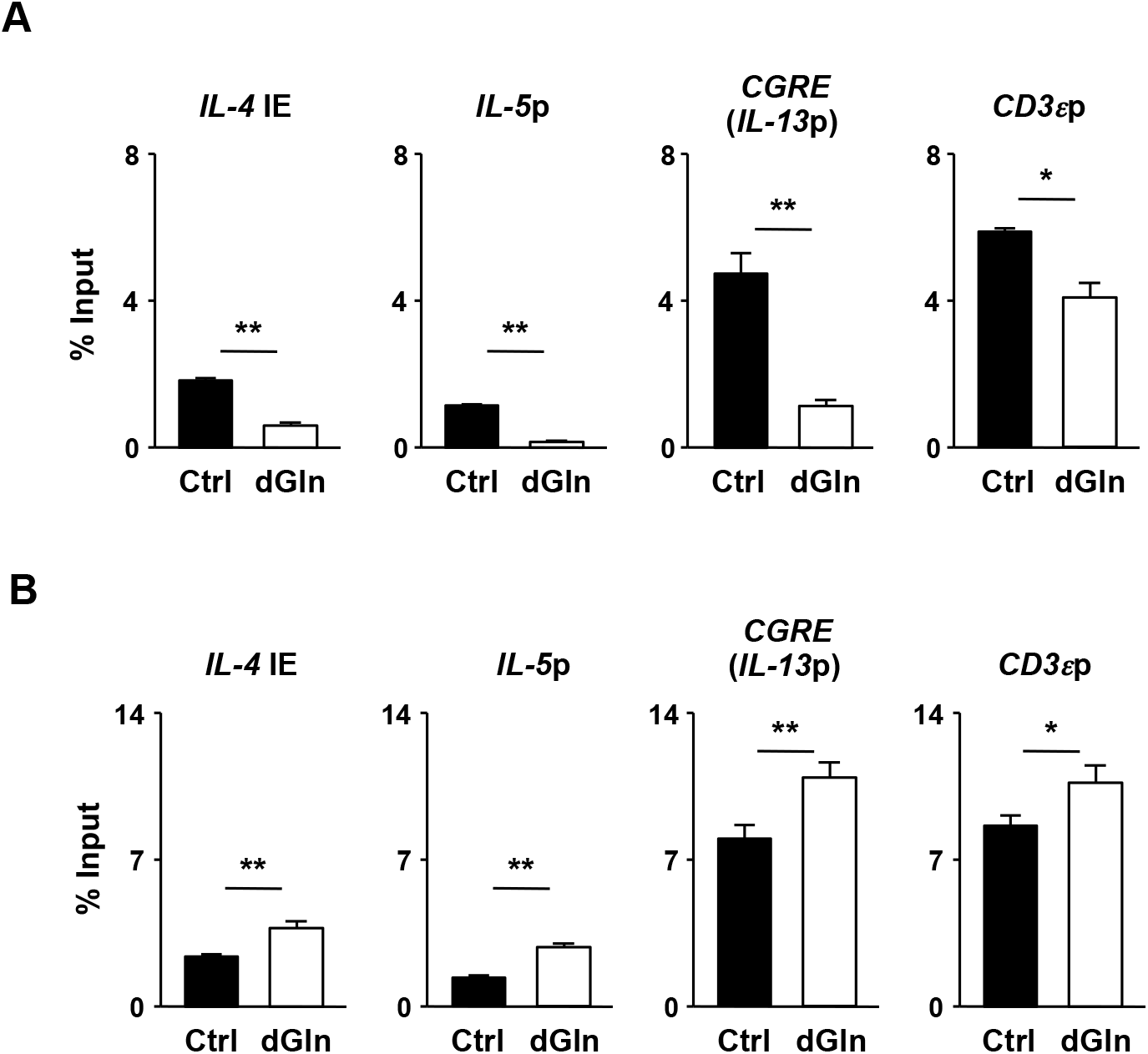
Histone modifications at the Th2 cytokine gene locus in CD4 T cells cultured under glutamine-deprived Th2 conditions. The levels of histone H3K9 acetylation (**A**) and H3K4 tri-methylation (**B**) at the Th2 cytokine and CD3ε gene loci in naïve CD4 T cells cultured under normal (Ctrl) or glutamine-deprived (dGln) Th2 conditions for two days. The results are shown with the standard deviations (n = 3: technical replicates). **P<0.01 (Student’s t-test).

**Figure 3-Figure Supplement 1.**
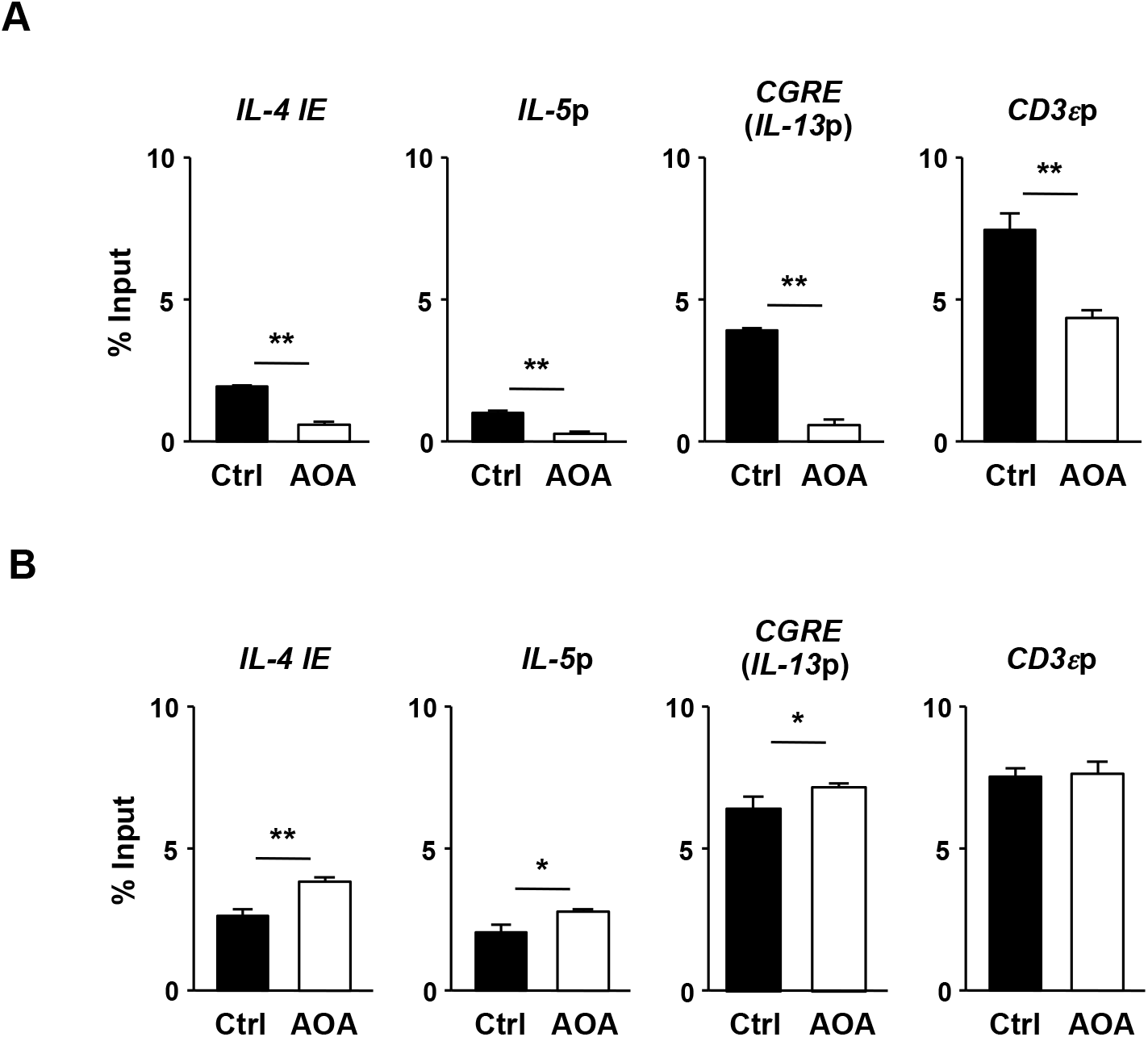
Effect of glutamine deprivation on histone modifications at the Th2 cytokine gene locus in CD4 T cells cultured under Th2 conditions. The levels of histone H3K9 acetylation (**A**) or H3K4 tri-methylation (**B**) at the Th2 cytokine and CD3ε gene loci in naïve CD4 T cells cultured under Th2 conditions in the presence (AOA) or absence (Ctrl) of AOA two days. The results are shown with the standard deviations (n = 3: technical replicates). **P<0.01 (Student’s *t*-test).

